# Ultrastructure of synaptic connectivity within sub-regions of the SCN revealed by genetically encoded EM tag and SBEM

**DOI:** 10.1101/2022.09.26.509467

**Authors:** Hugo Calligaro, Azarin Shoghi, Xinyue Chen, Keun-Young Kim, Yu Hsin Liu, Brian Khov, Benjamin Finander, Hiep Le, Mark H. Ellisman, Satchidananda Panda

**Affiliations:** Salk Institute for Biological Studies, La Jolla, CA, USA; Department of Neurosciences, University of California at San Diego School of Medicine, La Jolla, CA, USA; National Center for Microscopy and Imaging Research, University of California, San Diego, La Jolla, CA, USA

**Keywords:** melanopsin, Suprachiasmatic Nucleus, retinal ganglion cells, connectome

## Abstract

The suprachiasmatic nucleus (SCN) in the hypothalamus of the vertebrate brain is the central pacemaker regulating circadian rhythmicity throughout the body. The SCN receives photic information through melanopsin-expressing retinal ganglion cells (mRGC) to synchronize the body with environmental light cycles. Determining how these inputs fit into the network of synaptic connections on and between SCN neurons is key to impelling our understanding of the regulation of the circadian clock by light and unraveling the relevant local circuits within the SCN. To map these connections, we used a newly-developed Cre-dependant electron microscopy reporter, APEX2, to label mitochondria of mRGC axons, and serial blockface scanning electron microscopy to resolve the fine structure of mRGC in 3D volumes of the SCN. The maps thus created provide a first draft of the patterns of connectomic organization of SCN in the core and the shell, composed of different neuronal subtypes, and here shown to differ with regard to the patterning of their mRGC input as the shell receives denser mRGCs synaptic inputs compared to the core. This challenges the presently held view that photic information coming directly from the retina is mainly integrated by the core region of the SCN.

## INTRODUCTION

In mammals, circadian organization of physiology, behavior, and metabolism are necessary for a healthy lifespan. Although circadian rhythms are cell-autonomous and are found in almost every cell, the ventral hypothalamic brain region suprachiasmatic nucleus (SCN) localized above the optic chiasma plays an indispensable role in the circadian organization. Surgical ablation of the SCN or targeted genetic disruption of the circadian clock in the SCN abolishes behavioral circadian rhythm (Moore & Eichler, 1972; Stephan & Zucker, 1972). The neurons of the SCN generate autonomous circadian oscillations and orchestrate the timing of circadian rhythms in other tissues through synaptic and humoral regulation (Abrahamson & Moore, 2001; Welsh et al., 1995, 2010). Adaptation of circadian rhythm to seasonal changes in day length is also exclusively mediated through retinal light input to the SCN (Coomans et al., 2015). Given the central role of the SCN in the circadian organization and behavioral adaptation to ambient light, there is intense interest in understanding the cellular composition, cytoarchitecture, synaptic organization, and molecular properties of the SCN.

Each half of the SCN is composed of roughly 10,000 neurons and, based on their neuropeptide expression, is divided into two main subdivisions: the core (ventral) and the shell (dorsal). VIP- and GRP-expressing cells are located mainly in the core region, while AVP- and calretinin expressing cells are located mainly in the shell region of the SCN (Abrahamson & Moore, 2001; Fernandez et al., 2016; Moore, 1996; Yan & Silver, 2002). The SCN receives primary photic input from the retina and secondary photic input from the intergeniculate nucleus (IGL) and raphe nucleus (Abrahamson & Moore, 2001; Hannibal & Fahrenkrug, 2006; Welsh et al., 2010). The SCN neurons are believed to make extensive reciprocal connections among themselves and project to other hypothalamic and thalamic brain regions (Welsh et al., 2010). Another characteristic structure of the SCN is the presence of a communication network between neurons that does not depend on axonal communication but on another type of synapse, the chemical dendro-dendritic synapses (DDCS). These structures are relatively rare in the mammalian brain and their role has been little studied. DDCS have been identified in several types of peptidergic neurons of the hypothalamus (Alonso et al., 1985; F.-H. Güldner & Wolff, 1974; Theodosis et al., 1981; Theodosis & Poulain, 1984). These synapses have been proposed as serving to synchronize between neurons (Theodosis et al., 1981). In the SCN, the same function has been proposed, particularly between VIP neurons (Bosler & Beaudet, 1985) but, to our knowledge, the cell subtype possessing DDCS has not been identified to date. In a previous study, we established that SDCs are not isolated but form a network between cells in the SCN, and receive dense synaptic inputs from retinal and non-retinal sources (Kim et al., 2019). Their importance and distribution within the SCN remain however to establish.

While there is substantial progress in our knowledge of the gene expression pattern, electrophysiological properties, and peptidergic content of the SCN neurons, there is a limited understanding of the nature and extent of connectivity within the SCN. The SCN is composed of densely packed neurons with no apparent laminar or organizational stratification as seen in many other brain regions. Systematic sparse labeling of the rat SCN neurons and their cytoarchitecture has classified SCN neurons into subtypes based on the number of primary neurites (van den Pol, 1980). Electron microscopy image analyses of the SCN have also revealed a large variety of synaptic boutons of various origins, both outer-SCN (Card et al., 1981; Castel et al., 1993; Kim et al., 2019) and intra-SCN (Bosler & Beaudet, 1985; Castel & Morris, 2000), but their distribution and connectivity pattern remain largely unknown.

The emergence of connectomic EM methods such as serial blockface electron microscopy (SBEM) as well as genetically encoded electron microscopy tags, and viral-mediated expression of these tags in specific neurons has opened new avenues for investigation of the connectivity patterns of nuclei like the SCN. Recently, we have used a genetically encoded light and EM (GEM) tag (miniSOG) to specifically label the melanopsin expressing retinal ganglion cells (mRGCs) that monosynaptically connect to the SCN neurons and other brain regions (Kim et al., 2019). However, the nature of the GEM tag used in our prior study made it difficult to fully characterize the synaptic connectivity of the labeled cells and the study was not focused entirely on the SCN. We developed another tag system (Lam et al., 2014; Ramachandra et al., 2021) and modified this tag to allow viral-mediated expression in Cre-dependent mice in a way that enables specific labeling of the mitochondria, a strategy that can be done with mini-SOG or APEX2 to restrict marker densification to the mitochondria, leaving other subcellular domains with more native electron density. A similar approach was recently shown by Zhang and colleagues (Zhang et al., 2019). Using this label, we can identify and characterize the synaptic connectivity of the mRGCs and the rest of the connections within the SCN. Here we carried out a comparative analysis of the two main subdivisions of the SCN, the core and the shell, and interrogated: 1) What are the properties of the SCN cells (density, nuclei volume, number of neurites)? 2) What are the connectivity properties of the SCN neurons (somatic and dendritic synapses, balance between retinal and non-retinal input)? 3) What are the characteristics of axons and axonal boutons (volume, number of mitochondria, synaptic vesicles, dendritic intrusions, synaptic partners) and the specificities of mRGCs axons?

We found the core and the shell SCN differ in the composition of bipolar and multipolar neurons, associated with a small density of astrocytes. We confirmed the existence of a DDCS network and found around 10% of all dendrites are part of this network in the core but not in the shell. In both regions, DDCS-positive dendrites receive denser synaptic input from mRGCs and non-retinal axons. Finally, characterization of the retinal and non-retinal synaptic boutons revealed the mRGC boutons constitute a surprisingly larger fraction of boutons in the shell. However, the non-retinal boutons are larger, make more synaptic contacts, and can be divided into two groups depending on the presence of dendritic intrusions.

Together, these results bring us better insights into the complexity of SCN connectivity and the regional specialization of photic information integration. They also constitute a framework to understand the age and disease-dependent deterioration of circadian organization.

## RESULTS

### 1. SBEM of mouse SCN subregions receiving mRGC inputs labeled with genetically encoded EM tags

To comprehensively study the ultrastructure of the mouse SCN, we used a method established in our lab to intravitreally inject an adeno-associated viral vector expressing an EM reporter in presence of Cre-recombinase expressed in the melanopsin expressing retinal ganglion cells (Opn4^Cre/+^ mice; Kim et al., 2019). Unlike in our previous study that used a plasma membrane delimited EM reporter, mini-SOG, followed by photo-oxidation of the tissue block, we use a different EM reporter, APEX2 (AAV-EF1α-DIO-mito-V5-APEX2). We used a specific tag to address APEX2 to the mitochondria membrane to permit the recognition of mRGCs boutons in the SCN and conserve the axon membrane label-free to facilitate the identification of synapses (Figure 1). The use of this reporter does not require photo-oxidation, thus, is easier to stain and unlike membrane delimited mini-SOG, preserves the ultrastructural details of the labeled axons. These features allow large-scale staining (Lam et al., 2014; Martell et al., 2012) of neural tissues without toxicity.

We co-injected the EM reporter with a fluorescent microscopy compatible Cre-dependent reporter DIO-tdTomato (AAV-EF1α-DIO-TdTomato) into both eyes of Opn4^Cre/+^ mice. Four weeks post-injection, we verified the comprehensive labeling of mRGCs in both eyes and then processed the animal’s brain for SBEM following a published protocol (Lam et al., 2014).

Coronal brain sections of the hypothalamic regions were scanned for the SCN mid-way along the rostro-caudal axis. Then the SCN was divided into core, defined as the ventromedial SCN, and shell, the adjacent dorsolateral part. Each block was separately processed for SBEM. We collected 644 sections for the core and 537 sections for the shell with a thickness of 50 nm. We choose a section width of 50 nm to minimize the overlapping of small structures in the following sections such as synaptic vesicles that usually have a diameter of about 30 nm. We obtained a resolution of 7.4 nm per pixel.

**Figure 1.**
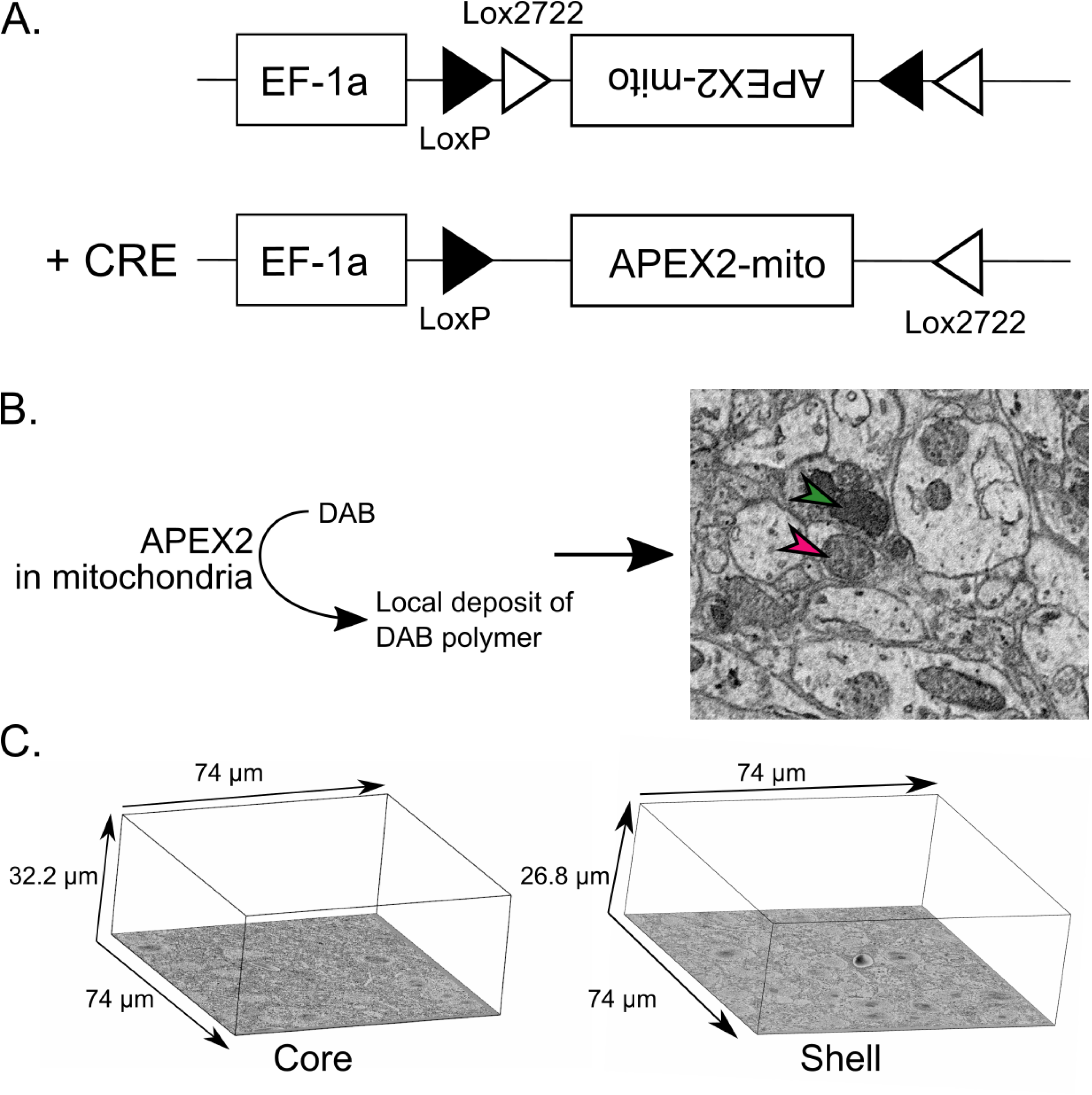
APEX2 specifically labels mRGCs mitochondria. **A.** Construction of the Cre-dependant APEX2 vector **B.** Enzymatic oxidation of DAB by APEX2 forms a local deposit of DAB polymer in mitochondria that appear dense to electrons in EM images. (red arrowhead = not labeled mitochondria, green arrowhead = labeled mitochondria). **C.** Representative image of the size of core and shell volumes.

In addition to the ultrastructural features of cells in the SCN, we also found axonal processes that contained darkly stained mitochondria, which are most likely from the mRGCs. As an initial random characterization of these axons with densely labeled mitochondria, we divided the volume on a 5 x 5 grid that resulted in 25 boxes. Out of the 25 boxes, 9 were selected and 5 to 10 axons per box containing darkly stained mitochondria were traced and segmented for their entire length contained within the volume. None of these axons traced back to a soma within the imaged volume, and were not myelinated, as expected for mRGC axons expressing the APEX2 tag. All of them contained more than one labeled mitochondrion. However, we found that among axons containing labeled mitochondria nearly 25.45% of axons were recognized and counted in the SCN shell and 22.92% of axons in the SCN core had at least one unlabeled mitochondrion. In other words, based on mitochondria staining in one section alone, we cannot conclusively classify an axon as being an mRGC or non-mRGC axon, and therefore, for all subsequent analyses, we traced any axon and checked for labeled mitochondria to classify them as mRGC axon. The unlabeled axons were likely of local origin constituting intra-SCN network or afferent projections from non-retinal sources.

### 2. Density and characteristics of SCN neurons in the core and the shell

In order to evaluate the cell characteristics in the core and shell of the SCN, we first attempted to segment all cells whose nuclei are visible in the respective volumes (Figure 2A). We found a slightly higher density of neurons in the shell compared to the core (core: 53.3 neurons per 100,000 µm³; shell = 67.5 per 100,000 µm³; Figure 2B). We traced and reconstructed the nuclei and computed their volume. We found relatively larger nuclei in the shell (core = 318.53 ± 7.35 µm^3^; shell = 346.45 ± 8.53 µm^3^; p<0.05; Figure 2C). We marked the astrocytes based on their characteristic features and found a similar density in both SCN regions (core: 6.7 per 100,000 µm^3^; shell: 5.7 per 100,000 µm^3^, Figure 2B). Too few astrocyte nuclei were fully inside the volumes for a meaningful comparison of their volume.

**Figure 2.**
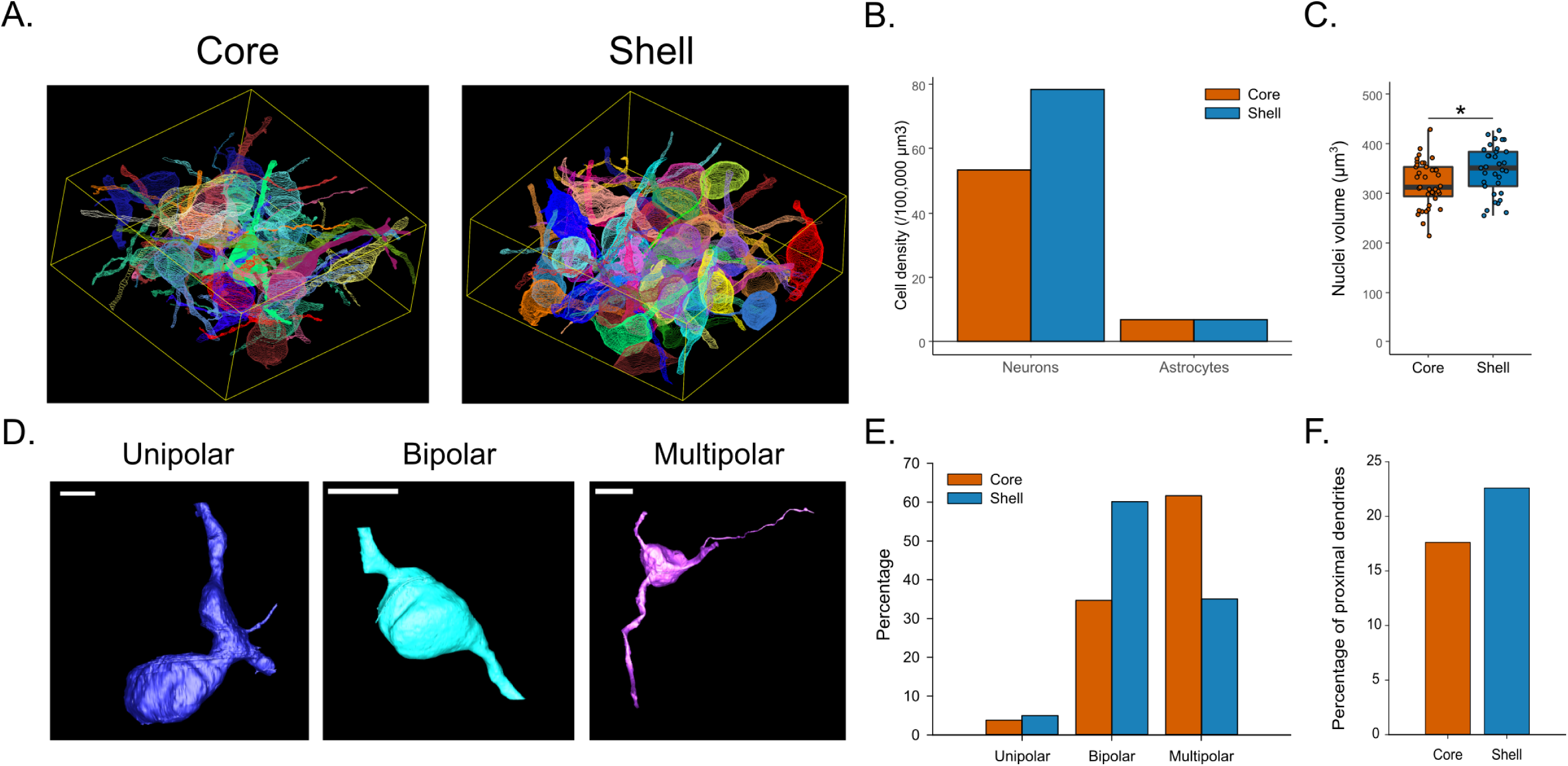
Density and types of SCN neurons. **A.** Representative image of 3D reconstruction of all complete neurons in the core (left) and the shell (right). **B.** Cell density of neurons and astrocytes. **C.** Volume of SCN neurons nuclei in the core and shell. D. Representative 3D reconstruction of unipolar, bipolar and multipolar neurons. **E.** Proportion of unipolar, bipolar and multipolar neurons in the core and the shell. **F.** Proportion of proximal dendrites in 100 randomly selected dendrites. *: p<0.05

We comprehensively segmented the processes of the neurons whose soma was fully inside the volume (core = 26 neurons; shell = 20 neurons). Based on the primary neurites extending from the soma, we classified neurons as unipolar, bipolar, or multipolar (Figure 2D). The unipolar neurons are scarce and accounted for <10% of each volume. Most SCN neurons are either bipolar or multipolar, indicating a regional specialization, with more bipolar neurons in the shell (core: 34.6%; shell: 60%) and more multipolar neurons in the core (core: 61.5%; shell: 35%, Figure 2E).

The number of dendrites extending from the SCN neurons reflects the amount of axodendritic input on a macroscale and the diversity with which the dendrites project to their own region compared to other regions. We randomly selected 200 dendrites, fully skeletonized them, and quantified which were proximal dendrites, defined as dendrites from a neuron whose soma is within our collected volume. Out of the dendrites we were able to fully skeletonize, 17.6% of dendrites are proximal dendrites in the core (31 dendrites out of 176 dendrites), while 22.5% of dendrites are proximal dendrites in the shell (40 dendrites out of 177 dendrites traced). This indicates that the shell has more proximal dendrites and thus that the shell neurons may form a more local network (Figure 2F).

To further characterize the neurons in both regions, we compared the density of nucleoli, a sign of high protein synthesis activity in the nucleus, and stigmoid bodies, a cytoplasmic structure specific to hypothalamic regions of the brain. We found a similar density of nucleoli in neurons in the core (1.44 ± 0.15 nucleoli/nuclei) and in the shell (1.15 ± 0.06 nucleoli/nuclei, p=0.219; Supp Figure 1A). In the core, there is an estimation of 10 stigmoid bodies per 100,000 µm³, and 28 stigmoid bodies per 100,000 µm³ in the shell (Supp Figure 1B).

### 3. SCN neurons in the core and the shell receive dense somatic and proximal synaptic input

As the core and the shell of the SCN contain different subtypes of neurons, we evaluated the connectivity of neurons’ soma and proximal axons to identify if one type of neuron receives more mRGCs synaptic input. Somas and proximal dendrites were fully segmented (core = 26 neurons; shell = 20 neurons). Then we systematically inspected the cell membrane of these neurons in all sections of the volumes to identify synapses. Neurites that presented the morphological characteristics of neurons (no post-synaptic synapses, small constant diameter) were excluded from the analyses. We found a total of 411 synapses in the core and 371 in the shell, contacting the soma or proximal dendrites (Figure 3A). We found a significantly higher density of synapses on soma in the shell (core = 8.38 ± 1.90 synapses; shell = 11.10 ± 1.56; p=0.026; Figure 3B) as well as for proximal dendrites (core = 13.65 ± 1.39 synapses/100µm; shell = 20.28 ± 174 synapses/100µm; p<0.001; Figure 3C).

**Figure 3.**
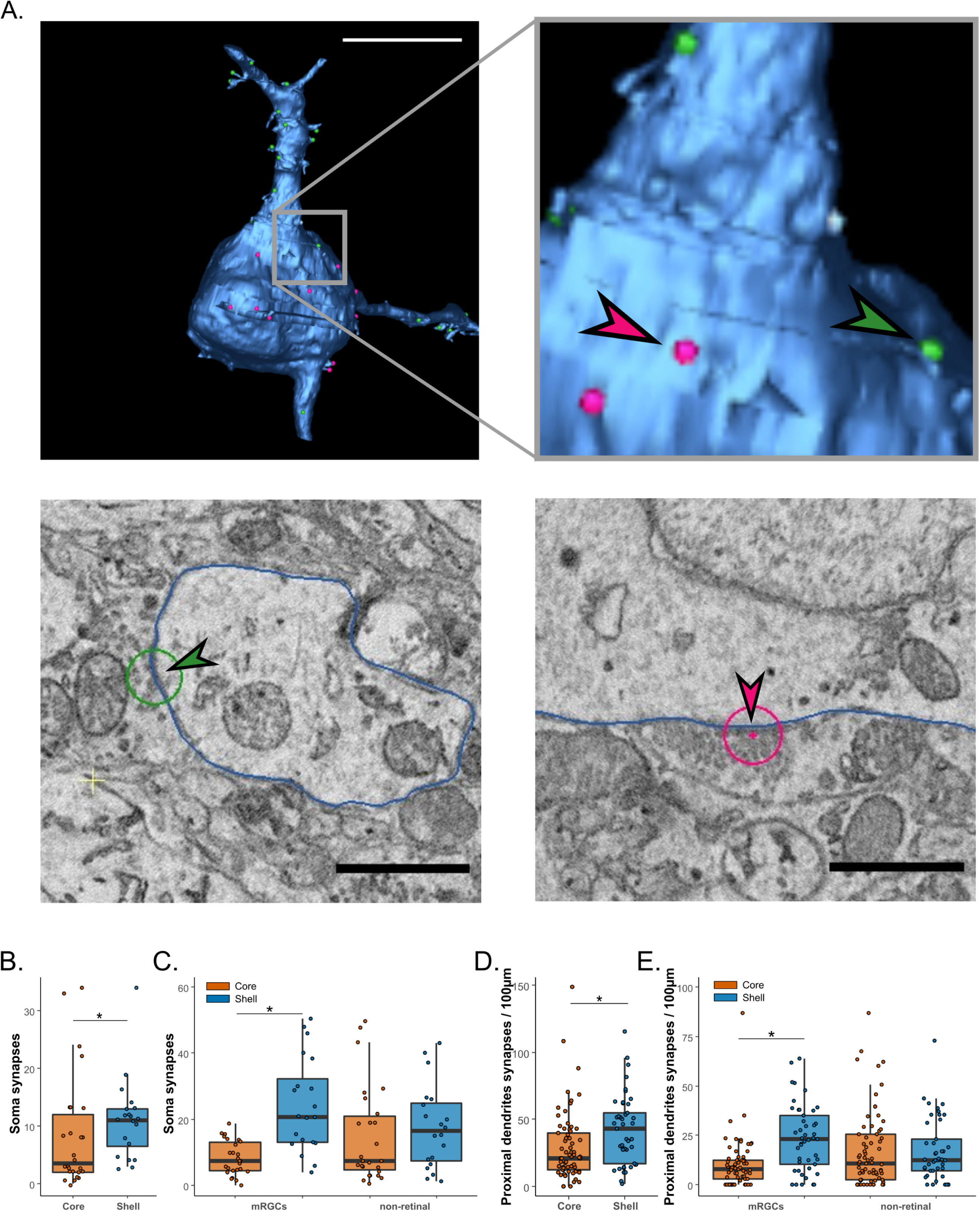
Proximal connectivity of SCN neurons. **A.** Representative image of 3D reconstruction of a neuron (upper panel) and EM images of synapses (lower panel) on proximal dendrite (green arrow, left) and on soma (red arrow, right). **B.** Total number of synapses contacting the soma. **C.** Total number of synapses contacting proximal dendrites. **D.** Number of mRGCs and non-retinal synapses contacting the soma. **E.** Number of mRGCs and non-retinal synapses contacting proximal dendrites. *: p<0.05

We then refined the analysis by using the APEX2 labeling to classify the synapses as being formed with mRGCs (with APEX2 labeled mitochondria) or non mRGCs axons. We found that mRGC synaptic density in the shell is significantly higher than in the core, on both soma (core = 3.58 ± 0.68 synapses; shell = 6.70 ± 0.72; p=0.003; Figure 3D) and proximal dendrites (core = 10.01± 1.48 synapses/100µm; shell = 23.52 ± 2.51 synapses/100µm; p<0.001; Figure 3E). On the contrary, we found no significant differences of the non-retinal synaptic density on the soma (core = 4.73 ± 1.44 synapses; shell = 4.05 ± 0.71; p=0.35; Figure 3D) or proximal dendrites (core = 17.29 ± 2.29 synapses/100µm; shell = 17.03 ± 2.34 synapses/100µm; p=0.595; Figure 3E).

Finally, we used the classification of neurons established earlier as bipolar or multipolar to verify if a particular subtype of neurons received more of one type of connection, but the synaptic density is homogeneous among all groups in both volumes with one exception; proximal dendrites of bipolar neurons receive more mRGCs synapses than multipolar neurons in the shell (Supp Figure 2A-D).

### 4. SCN neurons form a dense dendro-dendritic chemical synapse network

In addition to the direct synaptic contact on soma and proximal dendrites, we studied the density of axodendritic chemical synapses (ADCS) input on distal dendrites and the prevalence of dendro-dendritic chemical synapses (DDCS). We previously showed that DDCS are forming a network between SCN neurons (Kim et al., 2019).

To estimate the density of the DDCS network in the two regions of the SCN, we randomly selected 100 dendrites from the core and the shell volumes and searched for synapses where the associated element (pre- or post-synaptic) is also a dendrite (see Methods for the criteria of identification used; Figures 4A; Supp Figure 3).

In the core, 10.6% of dendrites present at least one DDCS while only 1.9% of dendrites in the shell form DDCS (Figure 4B). We found however a similar frequency of DDCS in DDCS-positive dendrites in the core and the shell (core = 2.60 ± 0.91 DDCS/100 µm, shell = 1.75 ± 0.22 DDCS/100 µm; p=0.79, Figure 4C). DDCS were identified previously in the SCN but we took advantage of SBEM to characterize further DDCS. Dendrites with DDCS do not show any morphological change close to the DDCS. The contact between the two dendrites is usually directly on the dendrite shaft. One occurrence of a DDCS on a dendritic spine has been observed in our volume. The DDCS are surrounded by smooth endoplasmic reticulum and at least one mitochondrion within 1 µm. The cluster of synaptic vesicles around the DDCS have similar densities in the core and shell (core = 375.34 ± 76,78 synaptic vesicles/µm^3^, shell = 593.26 ± 106.16 vesicles/µm^3^; p=0.15, Figure 4D). We noticed two types of DDCS, some have a very limited number of synaptic vesicles clustering on the synapse while others have a larger population of synaptic vesicles. However, the limited number of DDCS found in both volumes due to their low frequency prevents to further characterize them here. Future work on larger volumes might help establish if different types of DDCS exist within the SCN.

**Figure 4.**
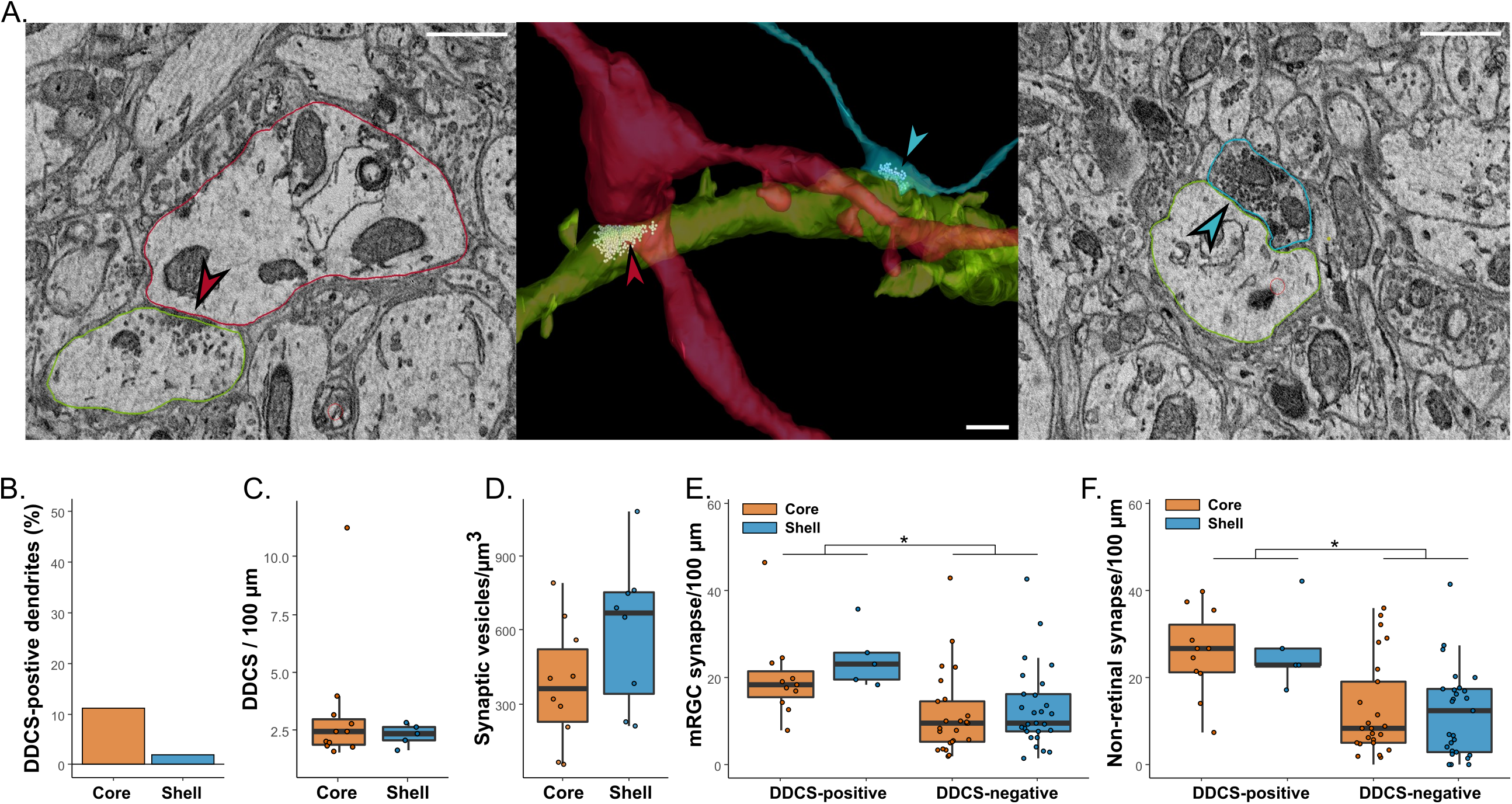
Network of dendro-dendritic synapses in the SCN. **A.** 3D reconstruction (middle) and representative image of EM image of DDCS (left, red arrow) and ADCS (right, blue arrow). **B.** Percentage of dendrites forming at least one DDCS. **C.** Frequency of DDCS along DDCS-positive dendrites. **D.** Synaptic vesicle density at the proximity of DDCS. E. Frequency of mRGCs synapses on DDCS-positive and negative dendrites. **F.** Frequency of non-retinal synapses on DDCS-positive and negative dendrites. *\*: p<0.05*

Among these DDCS-positive dendrites, we measured and determined the frequency of ADCS with mRGCs and non-retinal axons. The same analyses were conducted on a similar number of dendrites that don’t form DDCS (DDCS-negative dendrites). We counted a total of 720 synapses in the core and 501 in the shell. The DDCS positive dendrites receive a significantly higher number of synapses than the DDCS-negative dendrites in both the core and the shell (DDCS-positive: core = 53.24 ± 5.45 synapses/100 µm, shell = 57.08 ± 7.14 synapses/100 µm; DDCS-negative: core = 25.52 ± 3.23 synapses/100 µm, shell = 25.50 ± 2.89 synapses/100 µm, p<0.001; Figure Supp 2E), despite having a similar total length (DDCS-positive: core = 67.45 ± 6.25 µm, shell = 70.03 ± 8.84 µm, p=0.8269; DDCS-negative: core = 54.28 ± 2.94 µm, shell = 47.20 ± 2.95 µm, p=0.099; Figure Supp 2F). After classifying the synaptic boutons as mRGCs or non-retinal, we observed a similar pattern. The synaptic density is similar the core and the shell, but higher in the DDCS-positive dendrites compared to DDCS-negative dendrites for both mRGCs ADCS (DDCS-positive: core = 20.10 ± 2.97, shell = 24.46 ± 3.11; DDCS-negative: core=11.43 ± 1.90, shell = 12.93 ± 1.79, p<0.001; Figure 4E) and non-retinal ADCS (DDCS-positive: core = 25.75 ± 2.93, shell = 26.32 ±4.21; DDCS-negative: core=12.46 ± 2.24, shell = 11.82 ± 1.96, p<0.001; Figure 4F). A small fraction of boutons (∼5%) forming synapses with these dendrites were not identified as either mRGCs or non-retinal because they did not have mitochondria visible. These boutons were mostly close to the edge of the volume, reducing our capacity to follow the axon to find mitochondria allowing their identification.

In summary, dendrites forming a DDCS network receive similar synaptic input from mRGCs and non-retinal axons in the core and the shell and receive more synaptic input than DDCS-negative dendrites. DDCS are however more frequent in the core, suggesting a denser network of interconnected dendrites/neurons compared to the shell.

### 6. Myelinated and non-myelinated axons in the SCN

Two types of axons are present in the SCN: myelinated axons and non-myelinated axons that can participate in the SCN connectivity (Figure 5A).

We fully segmented myelinated axons in both volumes and examined their density, length, and branching pattern. None of them presented APEX-labeled mitochondria and were thus not originating from the retina. We identified 85 myelinated axons in the core and 54 in the shell, which correspond to a slightly higher density of myelinated axons in the core compared to the shell (core = 48.21 axons/100,000 µm³, shell = 33.62 axons/100,000 µm³; Figure 5B). The myelinated axons are significatively longer in the core compared to the shell (core = 43.10 ± 2.69, shell = 33.62 ± 2.97; p <0.01; Figure 5C), possibly due to the proximity of the optic chiasma. In the shell, none of the total 54 myelinated axons has branches while in the core, 4 out of the total 85 myelinated axons are branching once. In both volumes, only two axons showed demyelination but no boutons or synapses have been observed. These myelinated axons did not appear to participate in the SCN connectome of our two volumes. They might be connecting to an adjacent brain region or a different part of the SCN.

The non-myelinated axons, however, are highly participating in the SCN connectome. To analyze the characteristics of non-myelinated axons, we randomly selected 50 mRGCs and non-retinal axons in both SCN volumes, manually traced them, and quantified their length and the frequency of boutons (Figure 5D). None of the axons were traced back to the cell soma contained in our volume. We did not observe any difference in the average length of mRGCs or non-retinal axons (mRGCs: core = 42.26 ± 4.52 µm; shell = 40.79 ± 4.40 µm, p=0.7289; non-retinal: core = 43.51 ± 3.14 µm; shell = 36.66 ± 2.91 µm, p=0.108; Figure 5E) or the frequency of boutons along these axons (mRGCs: core = 9.45 ± 0.95 boutons/100μm; shell = 10.93 ±1.03 boutons/100 μm; p=0.1577; non-retinal: core = 11.45 ± 1.31 boutons/100μm; shell = 13.02 ± 1.42 boutons/100 μm; p=0.791; Figure 5F). The larger number of mRGCs contacts with DDCS-positive dendrites cannot thus be explained by a higher density of boutons or frequency of synapses. It rather suggests a synaptic preference of mRGCs for DDCS-positive dendrites in the core and the shell SCN.

**Figure 5.**
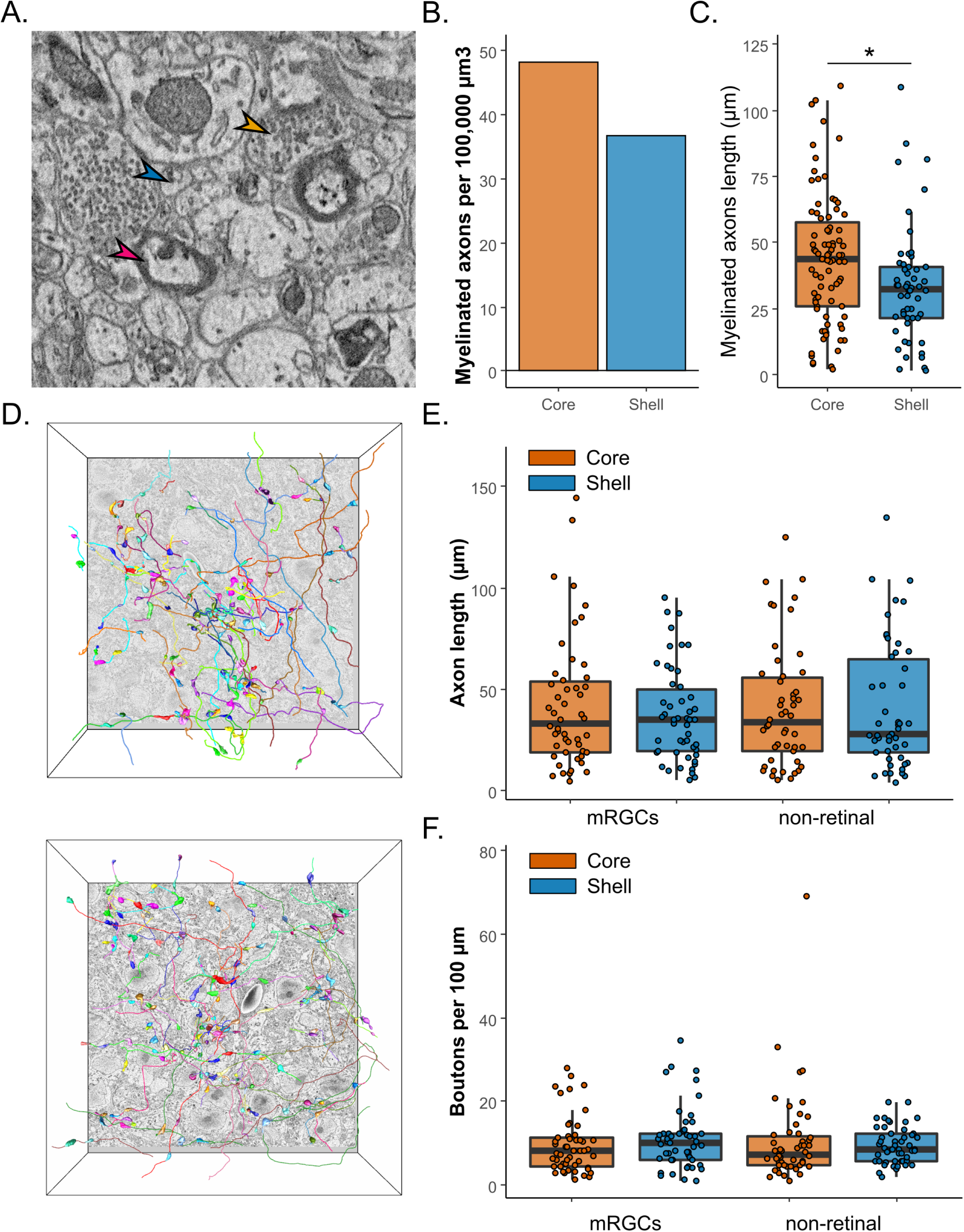
Myelinated and non-myelinated axons. **A.** Representative image of axons found in the SCN. Myelinated axons (red arrow), non-myelinated axons (blue arrow) and axonic boutons (orange arrow). **B.** Density of myelinated axons in the SCN volumes. **C.** Length of myelinated axons in the SCN. **D.** Representative image of the fully segmented axons and boutons in the core (upper panel) and the shell (lower panel). **E.** mRGCs and non-retinal axons length. **F.** Boutons frequency along mRGCs and non-retinal axons. *: p<0.05

### 5. mRGCs and non-retinal boutons have different morphological characteristics

#### a. Density and volume of mRGCs and non-retinal boutons

As we saw that synaptic input is not homogeneous in the SCN, we went further to characterize axonic boutons in the core and the shell. We selected 1 box out of the 5×5 grid previously created for both SCN volumes. In this box we exhaustively searched for all boutons and labeled them as mRGCs or non-retinal boutons, depending on the presence of labeled mitochondria (Supp Figure 3). A small fraction of boutons was not identified when they did not present mitochondria and were not connected to another bouton in the volume.

In the core, we counted a total of 675 boutons: 38.9% of mRGCs boutons, 55.5% of non-retinal, and 5.4% of unidentified boutons. In the shell, we counted a total of 601 boutons: 61.1% of mRGCs boutons, 31.4% of non-retinal boutons, and 7.4% of non-identified boutons. Based on the volume of the boxes (core SCN = 7052 µm³; dorsal SCN = 5880 µm³), we obtained a similar total density of boutons in both regions (core = 95.71 boutons/1000 µm³; shell = 102.19 boutons/1000 µm³). However, the density of mRGCs boutons is greater in the shell than in the core (core = 37.29 boutons/1,000 µm³, shell = 62.40 boutons/1,000 µm³; Figure 6A). On the contrary, the density of non-retinal boutons is greater in the core than in the shell (core = 37.29 boutons/1,000 µm³, shell = 62.40 boutons/1,000 µm³; Figure 6A).

**Figure 6.**
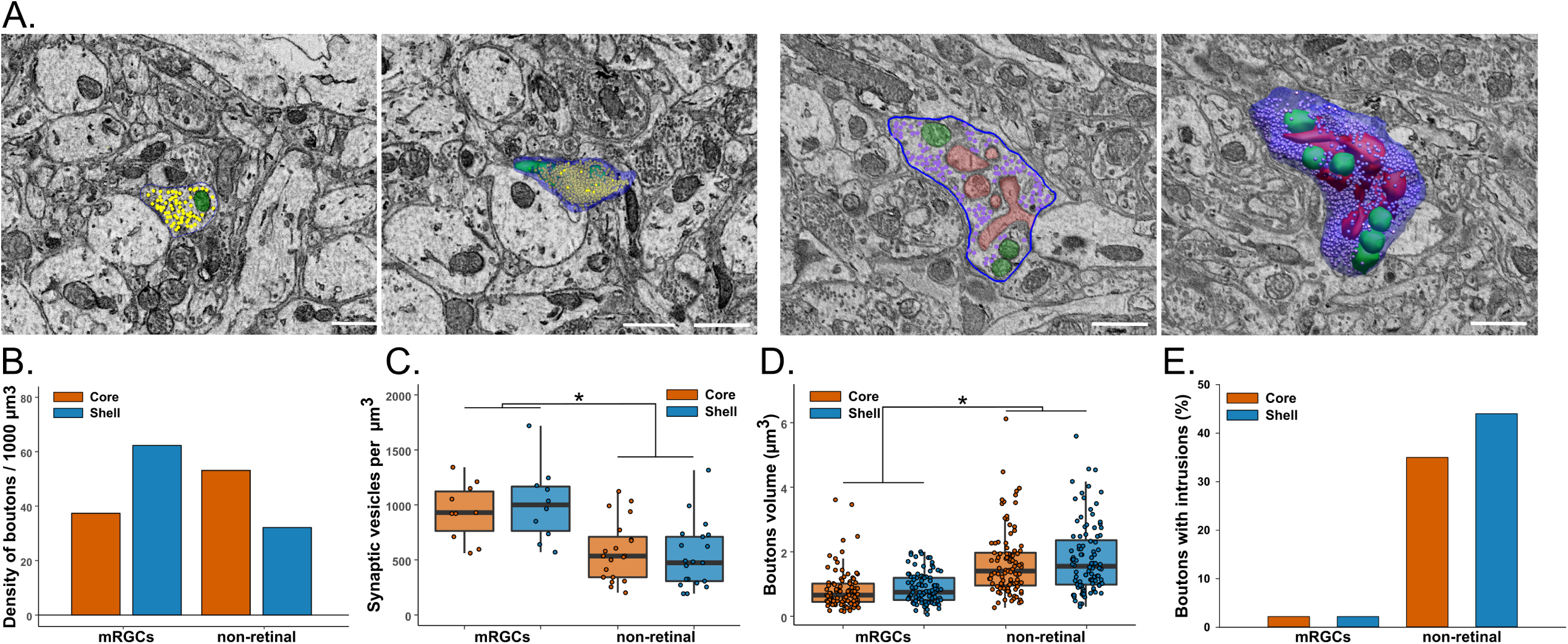
Characteristics of mRGCs and non-retinal boutons. **A.** Representative image of EM image and 3D reconstruction of boutons without (left) or with (right) intrusions. Green = mitochondria, red = dendritic intrusions, blue = cell membrane, dots = synaptic vesicles. **B.** Density of mRGCs and non-retinal boutons. **C.** Volume of mRGCs and non-retinal boutons. **D.** Density of synaptic vesicles in mRGCs and non-retinal boutons **E.** Percentage of mRGCs and non-retinal boutons containing intrusions. *: p<0.05

#### b. Characteristics of mRGC and non-mRGC boutons

We then randomly selected 100 mRGC and non-mRGC boutons and fully segment them and their content (Figure 6B). We did not observe a difference in volume of mRGC boutons (core = 0.83 ± 0.06 µm^3^; shell 0.88 ± 0.05 µm^3^; p = 0.12; Figure 6C) or non-retinal boutons (core = 1.66 ± 0.10 µm^3^; shell 1.80 ± 0.11 µm^3^; p = 0.40; Figure 6C) between the core and shell. However, non-retinal boutons are significantly larger than mRGCs boutons in both regions (p<0.001). mRGCs boutons have, however, denser synaptic vesicles than non-retinal boutons (mRGCs = 972.47 ± 66.23 synaptic vesicles/µm^3^; non-retinal = 554.68 ± 44.49 synaptic vesicles/µm^3^; p <0.001; Figure 6D). In addition, we noticed that only a small fraction of mRGCs boutons presented dendritic intrusions (∼2%), a structure that increases the contact surface between pre- and post-synaptic elements, while a significant percentage of non-retinal boutons present dendritic intrusions (core = 35%; shell = 44 %; Figure 6E).

Non-retinal boutons with dendritic intrusions are larger in both core and shell (p<0.001; Figure 7A), and contain more mitochondria (without intrusions = 2.66 ± 0.25 mitochondria, with intrusions = 4.09 ± 0.39 mitochondria; p<0.001; Figure 7B) of significantly higher volume (without intrusions: 0.060 ± 0.004 µm^3^, with intrusions: 0.061 ± 0.003 µm^3^; p=0.021; Figure 7C) and have lower density of synaptic vesicles (without intrusions: 737.46 ± 62.37 synaptic vesicles/µm^3^, with intrusions: 371.89 ± 26.81 synaptic vesicles/µm^3^; p<0.001; Figure 7D). However, the larger size of boutons with dendritic intrusions cannot be explained only by the additional mitochondria and the dendritic intrusions (Figure 7E). As expected, the number of synaptic partners is increased in boutons with intrusions (without intrusions: 1.54 ± 0.12 µm^3^, with intrusions: 2.17 ± 0.19 µm^3^; p<0.001; Figure 7F).

Non-retinal boutons can thus be separated into two different categories that could correspond to different origins of the axons; intra- and extra-SCN, for instance.

**Figure 7.**
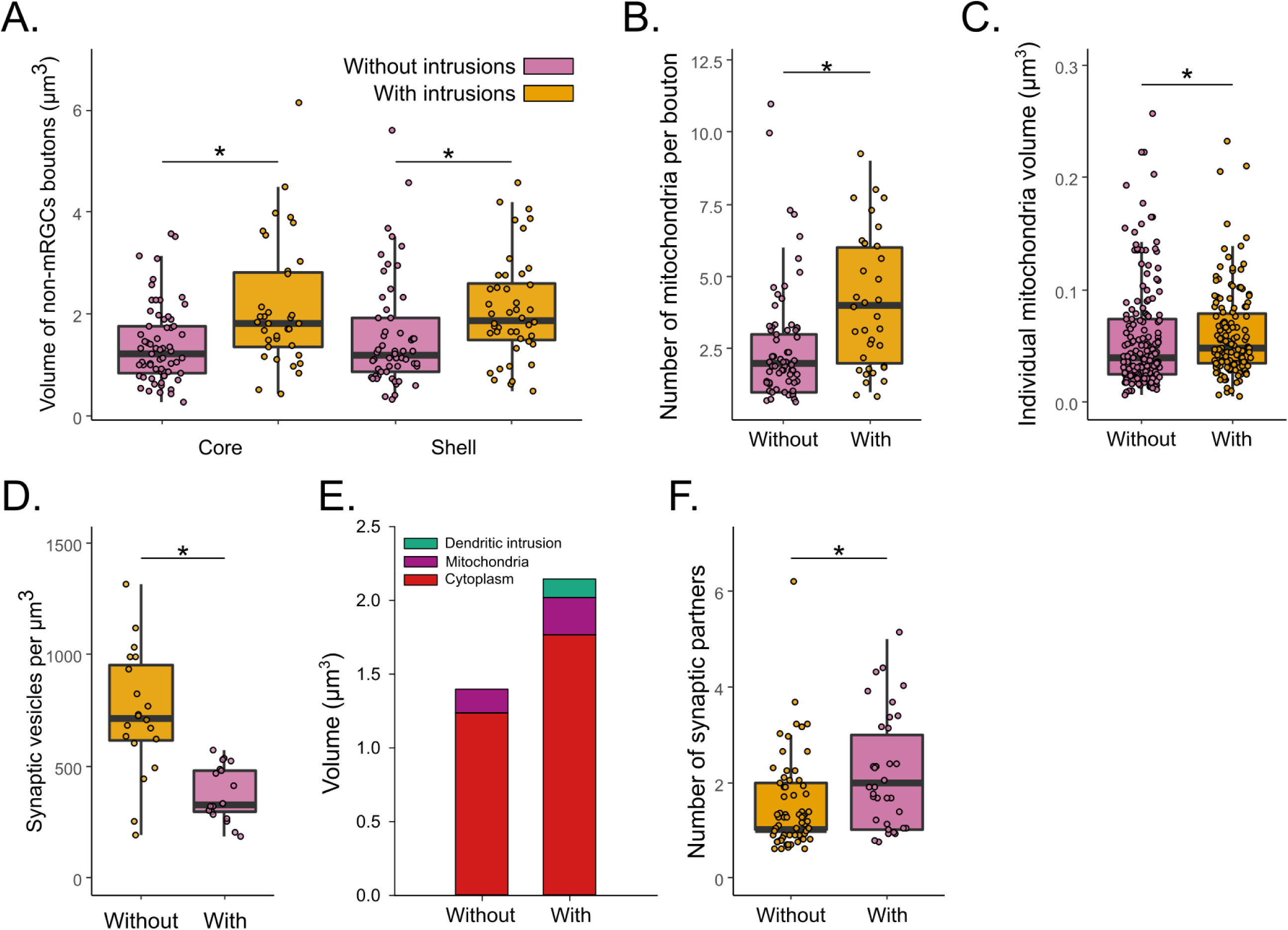
Characteristics of non-retinal boutons with dendritic intrusions. **A.** Volume of non-retinal boutons with and without intrusions. **B.** Number of mitochondria in non-retinal boutons with and without intrusions. **C.** Volume of individual mitochondria in non-retinal boutons with and without intrusions. **D.** Synaptic vesicle density in boutons with and without dendritic intrusions **E.** Volume proportion of non-retinal boutons with and without intrusions. **F.** Number of synaptic partners in non-retinal boutons with and without intrusions. *: p<0.05

## DISCUSSION

We presented here the results of the first inquiry of the SCN internal and external connectivity at this resolution. SBEM allowed us to access at an unprecedented scale how the retinal input is distributed in the two main subregions of the SCN, revealing the similarities and disparities.

### SCN neurons

Our results show that the shell region of the SCN is more densely packed with neurons than the core region. Previously, it was shown that the percentage of neurons in the rat SCN shell is 57% of all SCN neurons while the core contains 43% of SCN neurons (Moore et al., 2002). Our results support this observation, and it may explain the larger percentage of proximal dendrites found in the shell since more neurons are present in the shell. The density of astrocytes estimated from the current volume (∼70,000 astrocytes per mm^3^) is close to the number obtained from male rats SCN (68,627 astrocytes per mm^3^) (F. H. Güldner, 1983). The ratio of neurons/glial cells of about 10:1 is largely higher than in other brain regions. Glial cells have a crucial role in the rhythmic function of the SCN (Brancaccio et al., 2019). Unfortunately, the current volume contains too few glial cells to conduct a meaningful study of this type of cell in the SCN.

We classified SCN neurons as three subtypes based on their number of neurites (unipolar, bipolar, and multipolar), which is simplified from the classification introduced by van den Pol in 1980. van den Pol reported his discoveries on these rat SCN cell types: 1) a relatively small number of unipolar neurons; 2) the two dendrites coming from the bipolar neurons may branch once or twice, and these two dendrites are likely to change their direction, while the monopolar neurons do not; 3) radial multipolar neurons are found more commonly compared to other types, and most likely to be found in the dorsolateral SCN (the shell) (van den Pol, 1980). Our results partially match their finding. We observed a small number of unipolar cells, but most of the cells are bipolar or multipolar. However, we observed more multipolar in the ventral part than in the dorsal. These morphologically different neurons were then suggested to support specific electrophysiological functions (Jiang et al., 1997). The study found that bipolar cells had a higher membrane conductance and are likely to be innervated by RHT and be efferent cells also; the radial multipolar neurons are the only ones that did not respond with excitatory postsynaptic currents (EPSCs) to optic nerve stimulation and are speculated to be SCN interneurons. Again, our findings only partially confirm these hypotheses; we observed mRGCs synaptic inputs on all three types of neuronal cells (mono-, bi- and multipolar), either on the proximal dendrites or on the neuron soma. These morphological subtypes of SCN neurons could correspond to the different populations of neurons expressing specific neuropeptides. Initially, it was supposed that the neurons receiving the large majority of retinal input were VIP-expressing neurons (Lokshin et al., 2015). A subsequent study evaluated the retinal contact on the cell soma of the three main populations (VIP, AVP, GRP). They calculated that all cell types receive denser synaptic input from non-retinal origin (Fernandez et al., 2016). Our data show that retinal input corresponds to up to 40% of synaptic contacts on SCN neurons. Previous studies were done using light microscopy that has a lower resolution than in the present result and could have led to an underestimation of synaptic input on the soma by including synapses on proximal dendrites.

More interesting, we observed a specific higher density of mRGCs synapses contacts in the shell compared to the core. Fernandez and colleagues (Fernandez et al., 2016) showed that neurons from the shell require binocular light stimulation to induce C-Fos, which could explain why the synaptic density is almost doubled in the shell.

### Anatomy of boutons depends on axon origin

The SCN core mainly receives axonal projections from the intergeniculate leaflet (IGL) and the pretectal nuclei while the SCN shell receives inputs from limbic, hypothalamic, and brainstem nuclei (Abrahamson & Moore, 2001). In addition, both regions of the SCN receive photic information from the retina through mRGCs (Goz et al., 2008; Güler et al., 2008; Hatori et al., 2008; Hattar et al., 2006). The present study reveals for the first time the detailed architecture of mRGCs axons in the two subregions of the SCN. In addition to their difference in bouton volume, non-retinal and mRGCs boutons show various morphological specificities. Though both types of boutons contain a generic profile of synaptic organelles visible in EM images (mitochondria, vesicles, and ER), the main difference is the presence of dendritic intrusions. Significantly, close to half of the non-mRGC axons in multiple sets of samples contain intrusions of dendritic spines, while mRGCs boutons in the SCN hardly contain any. At the contact between these synaptic boutons and intrusions, some small synaptic vesicles were present.

Though it is mostly small vesicles surrounding the active zone, non-retinal boutons have much more diverse profiles of vesicles than mRGCs boutons. The mRGCs boutons mainly have small, packed vesicles and rare dense core vesicles accumulated at the presynaptic terminals, while non-retinal boutons could contain mainly big vesicles (∼150 µm), mainly small vesicles (∼40 nm), or a combination of big and small vesicles. According to previous literature on the origins of axons innervating the SCN, the big vesicles are possibly packed with serotonin (5-HT) from raphe nuclei afferents (Pelletier et al., 1981; Pickel & Chan, 1999) or neuropeptide Y (NPY) from geniculohypothalamic tract (GHT) (Card & Moore, 1982; Harrington et al., 1987; van den Pol, 2012), the small vesicles are possibly packed with γ-aminobutyric acid (GABA) from other SCN neurons or from GHT (Deken et al., 2003; Silver & Rainbow, 2015). In order to identify a synapse as excitatory or inhibitory, a gold standard in EM is to use specific features: excitatory synapses should have a dark post-synaptic density (“asymmetric synapse”), small round vesicles, and be preferentially on dendritic spines, while inhibitory synapses should not have a post-synaptic density (“symmetric synapse”), elongated synaptic vesicles and rarely on dendritic spines (Colonnier, 1968; Peters & Webster, 1991; Uchizono, 1965). We were not able to use these criteria in our dataset, as we were not able to clearly identify some elements. For example, we were not able to reproducibly identify post-synaptic densities, even on mRGCs synapses that are known to be in majority glutamatergic with a small population of GABAergic synapses (Castel et al., 1993; Purrier et al., 2014; Sonoda et al., 2020). We, therefore, decided not to use this classification in this study. Nonetheless, even without this characterization, our data still provide useful inside into the connectivity of the SCN.

The comparison of mRGC bouton volumes between ventral and dorsal SCN reveals the synaptic strength between mRGCs and the distinct parts of SCN. The synaptic strength can be indicated by both the current strength and the subsynaptic structures, such as the number of mitochondria near the synaptic cleft (Verstreken et al., 2005), synaptic vesicles (Karunanithi et al., 2002), bouton size/volume and more (Meyer et al., 2014). Our data showed that the shell has slightly denser boutons than the core SCN while the average counts of mRGC boutons on one axon are not significantly different between ventral and dorsal SCN, given that the axonal lengths and branching patterns are similar. Together, these data showed an equally dense mRGC projection into both regions of the SCN and a stronger synaptic connection between mRGC and the shell SCN which should still be evaluated further. Agreeing with our observation, early studies also show direct retinal innervation of both the ventral and dorsal SCN (Beier et al., 2020; Hattar et al., 2006). On the contrary, it was reported that the ventral SCN was reported to receive most of the retinal input and transfer this input to the dorsal SCN (Abrahamson & Moore, 2001; Kriegsfeld et al., 2004; Moore et al., 2002; Welsh et al., 2010).

Beyond the comparison between mRGCs axons and non-retinal axons, different features of the non-retinal axons bring a great discovery on a diversity of afferents innervating the SCN. Here, we used intrusions of dendritic spines to characterize boutons of axons with or without dendritic intrusions, previously associated with different functions (driver vs modulator of the neuronal activity; (Sherman & Guillery, 1998). The more intrusions of dendritic spines are in one bouton, the bigger this bouton is, and the more synaptic partners and mitochondria this bouton will have. This could reflect that having a larger population of mitochondria can support more synaptic activity and expand the volume of the bouton. The presence of intrusions of dendritic also suggests a difference in function and/or origin. As we showed that almost none of the mRGCs boutons received dendritic intrusions, we could hypothesize that axons originating from outside the SCN would form small boutons working as modulators, while axons of SCN neurons would form these boutons with dendritic intrusions working as driver during intra-SCN communication. Previous anatomical studies of the synaptic afferent of the SCN did not show dendritic intrusions in glutamatergic or serotoninergic axonic boutons (Bosler & Beaudet, 1985; Castel et al., 1993; Kiss et al., 1984). Intrusions-like patterns have been found in GABAergic boutons that don’t express AVP, probably originating within the SCN (Castel & Morris, 2000), but no intrusions were observed after specific staining of VIP cells (Card et al., 1981). However, as the main objective of these studies was not to identify boutons with intrusions, it is difficult to conclude further on which type of boutons contain dendritic intrusions. Further studies will be needed to clarify their origin and roles.

### Dendro-dendritic chemical synapses form a dense network in the SCN

Dendro-dendritic chemical synapses were first reported in the SCN via ultrastructural studies in the 1970s (F.-H. Güldner & Wolff, 1974) and were observed to be uncommon (Moore & Bernstein, 1989). Prior to that, the discovery of this type of synapse between mitral and granule cells in the mammalian olfactory bulb started a growing discussion of the function of DDCS (Rall et al., 1966; Shepherd, 2009). Even though extensive work was done to study DDCS in the olfactory system, the DDCS in SCN was barely investigated since it was found.

In a previous study, we observed that dendrites with DDCS form a network in the core SCN (Kim et al., 2019). We confirmed the presence of a dense network of DDCS-positive dendrites is present in the core and, although less dense, in the shell. The limited size of the volume studied here prevents us to determine if these two networks are interconnected or are functioning in parallel, with communication through regular ADCS. The homogeneous synaptic input from both mRGCs and non-retinal axons argues in favor of a continuous network. However, previous studies have shown that the core and the shell can be desynchronized during experiments such as jet lag (Albus et al., 2005; Davidson et al., 2009; Mei et al., 2018; Nakamura et al., 2005; Yamaguchi et al., 2013). In these conditions, the shell shows a rapid shift while the core shift more slowly, suggesting a separated process for entrainment to the new light cycle.

Here, we noticed two patterns of these DDCS, some have a very limited number of synaptic vesicles clustering on the synapse while others have a larger population of synaptic vesicles. Unfortunately, the present volume does not allow us to further characterize these synapses. However, given that most of the core SCN neurons are expressing GABA and VIP, we can suppose they are probably releasing either or a combination of them. The precise role of DDCS in the SCN remains to be elucidated but we can hypothesize a role in the communication between adjacent cells, as we fail to identify axo-dendritic or axo-somatic synapses from a proximal axon in either volume.

This communication could be crucial for the synchronization between SCN cells. Multiple pieces of evidence showed that extracellular GABA is rhythmic (Hastings et al., 2018) and drives synchrony between SCN cells. This paracrine GABA could be originating from the DDCS rather than axonal synaptic terminals. This hypothesis fits with recent results showing a form of a small-world synaptic network composed of interconnected nodes mainly in the core SCN, allowing a faster re-synchrony of neurons after tetrodotoxin treatment (Abel et al., 2016). In addition, DDCS-positive dendrites receive more retinal and non-retinal input, these GABAergic synapses could modulate their responses in a phase-dependent manner. Further studies are needed to elucidate the precise role of this DDCS network in the SCN and as a synaptic input integrator.

### SCN neurons contain specific intracellular organelles

In addition to the connectivity variation between both regions of the SCN, this dataset allowed us to identify specific ultrastructures that cannot be observed otherwise. In particular, we were able to identify the presence of stigmoid bodies (SB) in the mouse SCN, with a higher density in the shell compared to the core. SB was described as a structure similar to the nucleolus located in the cytoplasm of neurons. While its exact function still remains unclear, it was speculated to be involved in the aromatization of androgens to estrogens, since the use of the antibody against the placental antigen associated with aromatase P-450 reveals that the SBs are located in sex-steroid-sensitive peripheral tissues such as ovary and testis (Santolaya, 1973; Shinoda et al., 1993). These neurons are in multiple brain regions including the hypothalamus, thalamus, amygdala, septum, hippocampus, colliculi, and brainstem (Santolaya, 1973).

It is worthwhile to note that the different distribution of stigmoid bodies possibly reveals the different functions of the core and shell SCN in the content of the circadian rhythm. Multiple studies have previously identified dot-like structures in rodents’ brains that are 5-hydroxytrptamine_7_ (5-HT_7_) receptor immunoreactive (Muneoka & Takigawa, 2003). These structures match the size and location of the SB we observed in the present study. The expression of 5-HT_7_ was also found to be in the mice SCN based on the positive immunoreactivity (Belenky & Pickard, 2001). Though linking the existence of the stigmoid bodies and serotonergic neurons could bring an interesting discussion into the content of the circadian regulation, this link is not supported thoroughly since the EM examination of the location of 5-HT_7_ did not report the localization of 5-HT_7_ on the stigmoid bodies (Belenky & Pickard, 2001).

We observed a relatively low number of myelinated axons compared to unmyelinated ones. They appear to be passing through the SCN and made no synaptic connections with the local dendrites or somas in our volume, let alone terminate in it. The core has a higher density of myelinated axons than the shell, matching previous observations (van den Pol, 1980). A few mitochondria of these myelinated axons appeared darker, reminding of APEX2-labeled mitochondria. However, as the majority of other mitochondria of the axons were clearly not labeled, we attributed the darker coloration to visual contrasts of the EM image. Moreover, previous experimental evidence suggests that mRGCs axons are unmyelinated because of a low conduction velocity, compared to conventional RGCs (Cahill & Menaker, 1989; Do et al., 2010; F. H. Güldner, 1978; Shibata et al., 1984). In addition, we first reported here the demyelination of axon branches in the SCN. We can speculate that these branches are preparing to reach their synaptic targets. However, these branches were not observed to be synaptically active in the current volume. The possibility of abnormal demyelination is out of the reach of this research due to the absence of a temporal resolution but would be worth investigating in aging and neurodegenerative disease mouse models.

In summary, we established, for the first time, the mRGCs connectivity in two distinct regions of the SCN, the core and the shell, in a healthy mouse. We uncovered a multi-level system of regulation by light of the SCN activity. This work opens the possibility of studying the impact of neurodegenerative disease and circadian-related syndromes at the level of neuron connectivity.

## Materials and Methods

### Mice

All animal experiments and use were approved by the Salk Institute IACUC. 8-months old mice were used for protocol optimization. All mice were housed under 12h/12h Light / Dark cycles of 100 lux of white light, in a temperature-controlled room with food *ad libitum*. Collection of the SCN was done at ZT6. Both SCN volumes are from the same mouse.

### Vector construction

A mitochondria-specific sequence was cloned into the 3’ end of the APEX2 construct (Lam et al., 2014; Ramachandra et al., 2021) and was inserted in an inverted orientation between the lox sites in an AAV2-DIO vector (Cardin et al., 2009) to create AAV-DIO-Mito-V5-APEX2 (Addgene plasmid # 72480; RRID:Addgene_72480). To enable an extended search of the ROI under a fluorescent microscope, we co-injected the APEX2 virus with a tdTomato-farnesyl virus to the Opn4^Cre/+^ mice. AAV-DIO-Mito-V5-APEX2 and AAV2-DIO-tdTomato-farnesyl were produced by the Salk Gene Transfer, Targeting and Therapeutics Viral Vector Core Facility at titers of 2.0×10^11^ TU/ml and 9.4×10^11^ TU/ml, respectively.

### Vector Injection

Anesthesia was induced with isoflurane (4%) and maintained (2%) until the end of the procedure. The mouse is placed under a dissection microscope so one eye is completely visible. Gentle pressure is applied around the eye so the edge of the sclera is visible. A small incision is made with a 27-gauge insulin needle 0.5 mm posterior to the *Ora Serrata*. The vector is loaded into a Hamilton microliter syringe with a 34-gauge beveled needle mounted on a micromanipulator. The micromanipulator is used to insert the loaded needle through the incision. The vector is slowly injected and allowed to diffuse through the vitreous humor for 2 minutes. The whole procedure is then repeated on the other side with the appropriate vector. After both eyes have been injected, the animal is removed from isoflurane anesthesia and lubricant eye ointment (AKORN) is applied to both eyes. The animal is placed in a clean cage to recover and is returned to its home cage when righting reflex is restored, after 1-2 minutes.

### SBEM Staining and Imaging

Tissue was prepared for SBEM as previously described (Deerinck et al., 2010). The retina and brain volumes were collected in 2.0 to 2.4 kV accelerating voltages, with a raster size of 20k×20k or 24kx24k and pixel dwell time of 0.5 −1.5 μs. The pixel size was 7.4 nm and section thickness was 50 nm. Before each volume was collected, a low magnification (∼500×) image was collected of the block face to confirm the anatomical location of the volume based on tissue landmarks. Once a volume was collected, the histograms for the slices throughout the volume stack were normalized to correct for drift in image intensity during acquisition. Digital micrograph files were normalized using Digital Micrograph and then converted to MRC format. The stacks were converted to 8-bit, mosaics were stitched, and volumes were manually traced for reconstruction and analysis.

### Three-dimensional reconstruction and analysis

To analyze these datasets, we used the publicly available software package IMOD, specifically developed for the visualization and analysis of EM datasets in three dimensions ((Kremer et al., 1996); http://bio3d.colorado.edu/imod/). Cross-sectional contours were manually traced for consecutive data slices in the z dimension to determine the boundaries of user-defined objects. For some objects, contours were traced in every other data slice in the z dimension. These contour profiles were used for three-dimensional volumetric reconstruction of the cell body, axons, boutons, and organelles.

### Manual segmentation

#### Length and volume quantification

To measure the length of neurites, we fully traced each branch individually by marking the neurite center every 5 z-steps. To calculate the volume of organelles, we fully segmented them individually every 2 z-steps. The length and volume values are then extracted using *imodinfo*.

#### Synaptic input quantification

Synaptic sites were identified by scanning the neurites previously skeletonized and finding areas matching at least two of the following criteria: the presence of synaptic vesicles close to the cell membrane, proximity between the pre- and post-synaptic element cell membranes, and presence of dendritic intrusions. Synapses are then classified between ADCS and DDCS by identifying both elements using the specific criteria listed below. For ADCS, the axonic element is classified as either mRGCs or non-retinal based on the APEX2 labeling of mitochondria. To avoid over- and underestimation of the density of synapses on dendrites, we excluded from the analysis all dendrites shorter than 2 µm. This criterion excluded 3 proximal dendrites in the core.

#### Identification of axons and dendrites

To discriminate between axons and dendrites, we look for different characteristic elements. The dendrites have usually a consistent and large diameter, with dendritic spines, dendritic intrusions, and are mostly postsynaptic elements of synapses. The axons have a smaller diameter with swelling forming synapses, receive dendritic intrusions, and are systematically the presynaptic element of synapses.

#### Counting of nuclei, astrocytes, stigmoid bodies, nucleoli

To count the nuclei, astrocytes, stigmoid bodies, and nucleoli in the two data sets, a 5×5 grid was created. The entirety of the data set was scanned systematically by starting from one corner and going through all the created grids and marking each of the nuclei, astrocytes, stigmoid bodies, and nucleoli with a different color. This process was done until all the areas in the volume sets of the ventral and dorsal SCN were covered.

#### Bouton identification

For the identification of boutons, we looked for areas of the axon that were swollen to a diameter at least twice as large as the average diameter of the axon. Our criteria for swelling to be considered a synaptic bouton included at least two of the following: the presence of a post-synaptic density in the cell directly contacting the swelling, presence of at least one mitochondrion, evidence of synaptic vesicles close to the cell membrane, or presence of dendritic intrusions.

#### Identification of myelinated axons

For the segmentation of myelinated axons, we created a 5×5 grid and started at the lowest z-step in the data set, and looked for axons with thick dark myelin around them. These axons were traced to obtain an estimate of their length and branching pattern. This process was done until all the areas in the volume sets of the ventral and dorsal SCN were covered.

### Statistical analysis

Results are expressed as means ± standard error of the mean (SEM). All statistical analyses were performed using R (R project). Comparisons of the two groups were done using a non-parametric rank-sum test (Mann-Whitney U Test). Comparisons of multiple groups were done by using ANOVA on ranks (Kruskal-Wallis H test) followed by a post hoc test (Pairwise Mann-Whitney U-tests). A statistically significant difference was assumed with a p-value inferior to 0.05.

## Supplementary Figures

**Supplementary Figure 1.**
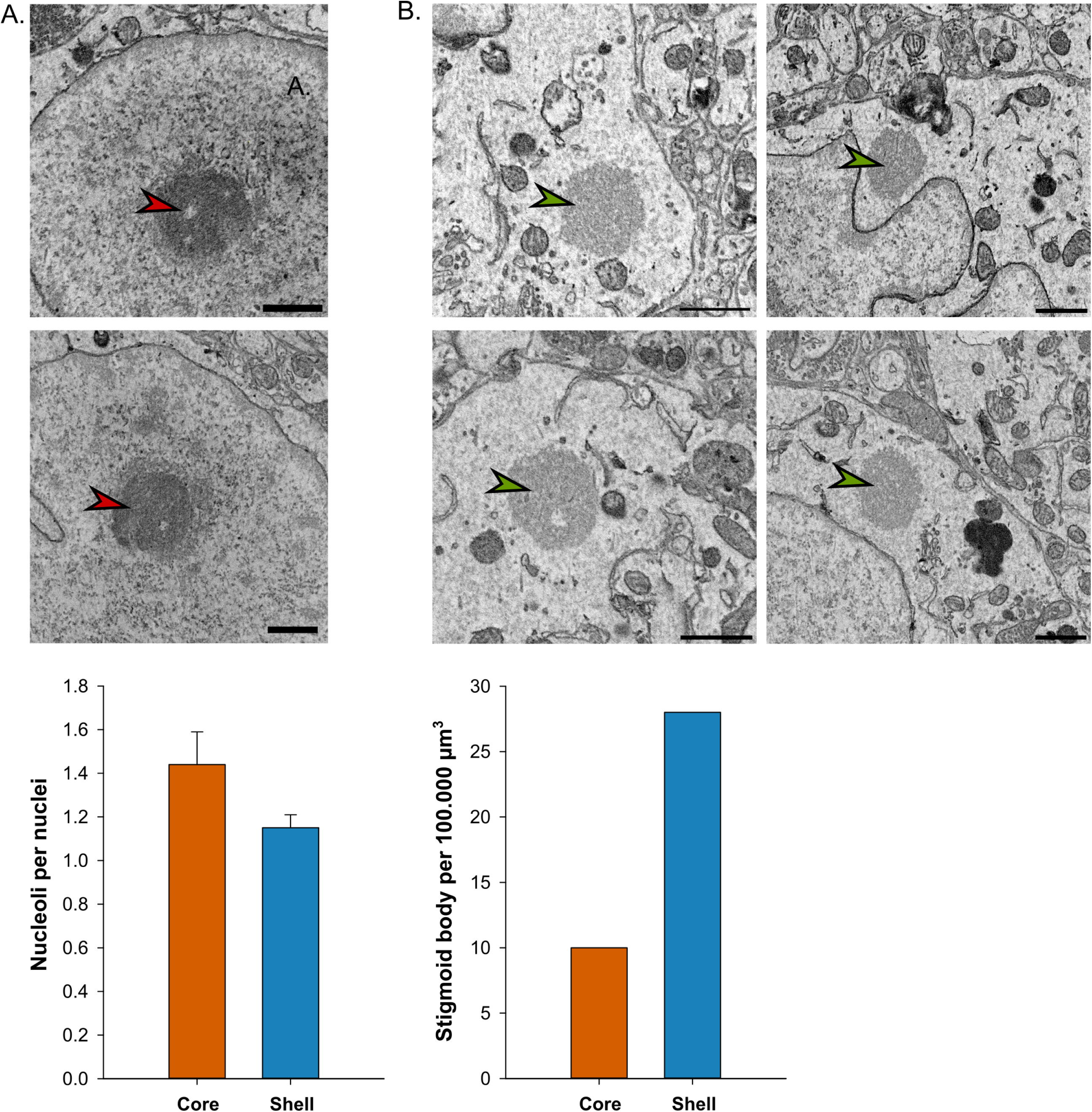
Nucleoli and stigmoid bodies in the Core and the Shell regions of the SCN. **A**. Representative EM image (red arrows upper panel) and frequency of nucleoli per nuclei (lower panel) in the core and the shell. **B.** Representative EM image (green arrows, upper panel) and density of stigmoid bodies (lower panel) in the core and the shell.

**Supplementary Figure 2.**
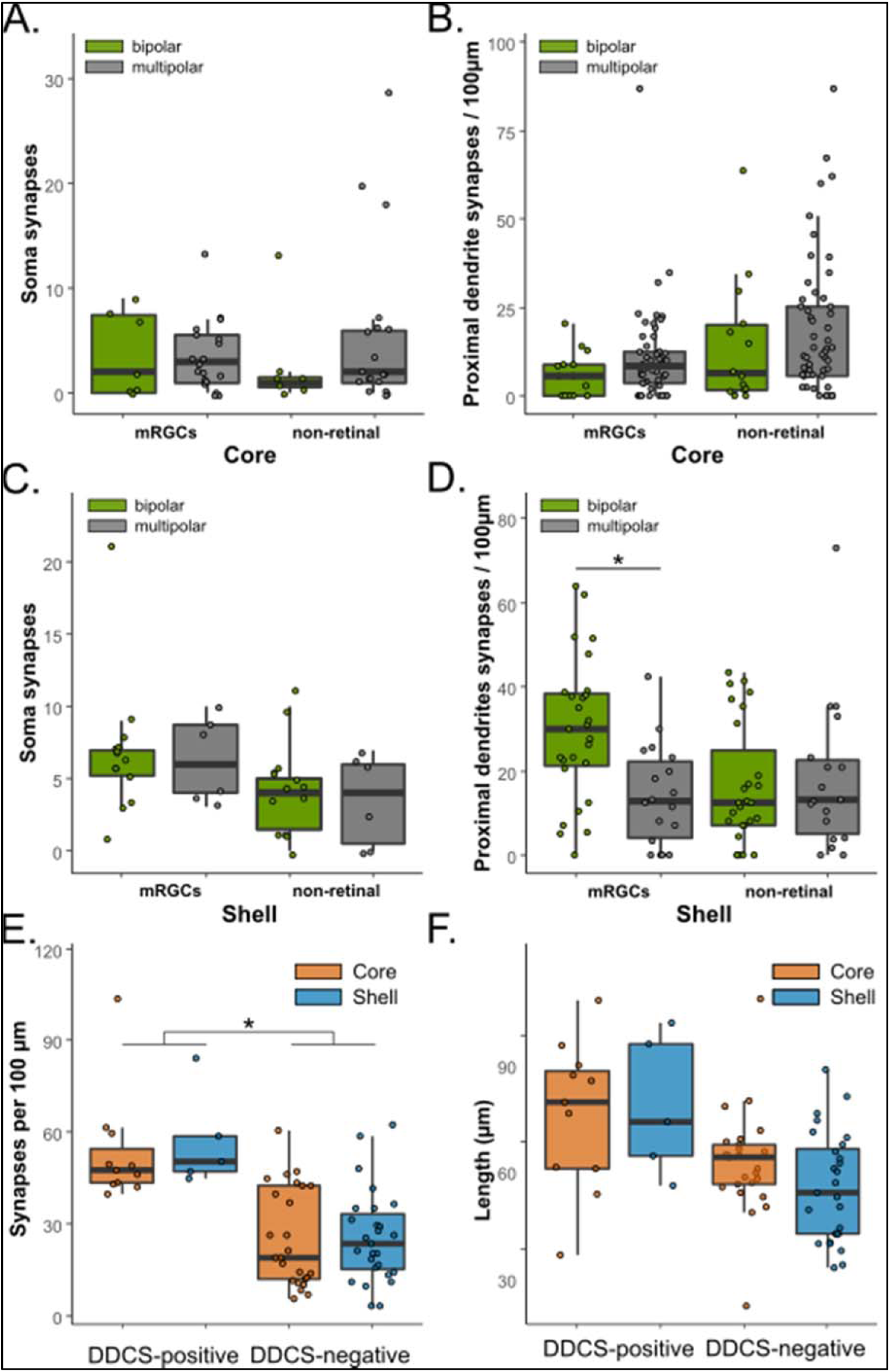
Synaptic densities on soma, proximal dendrites and distal dendrites. **A,C.** Synapse density on soma of bipolar and multipolar neurons in the core (A) and the Shell (C). **B,D.** Synapse density on proximal dendrites in the core (B) and the Shell (D). **E.** Frequency of ADCS on DDCS-positive and negative dendrites. **F.** Average length of DDCS-positive and negative dendrites. *: p<0.05

**Supplementary Figure 3.**
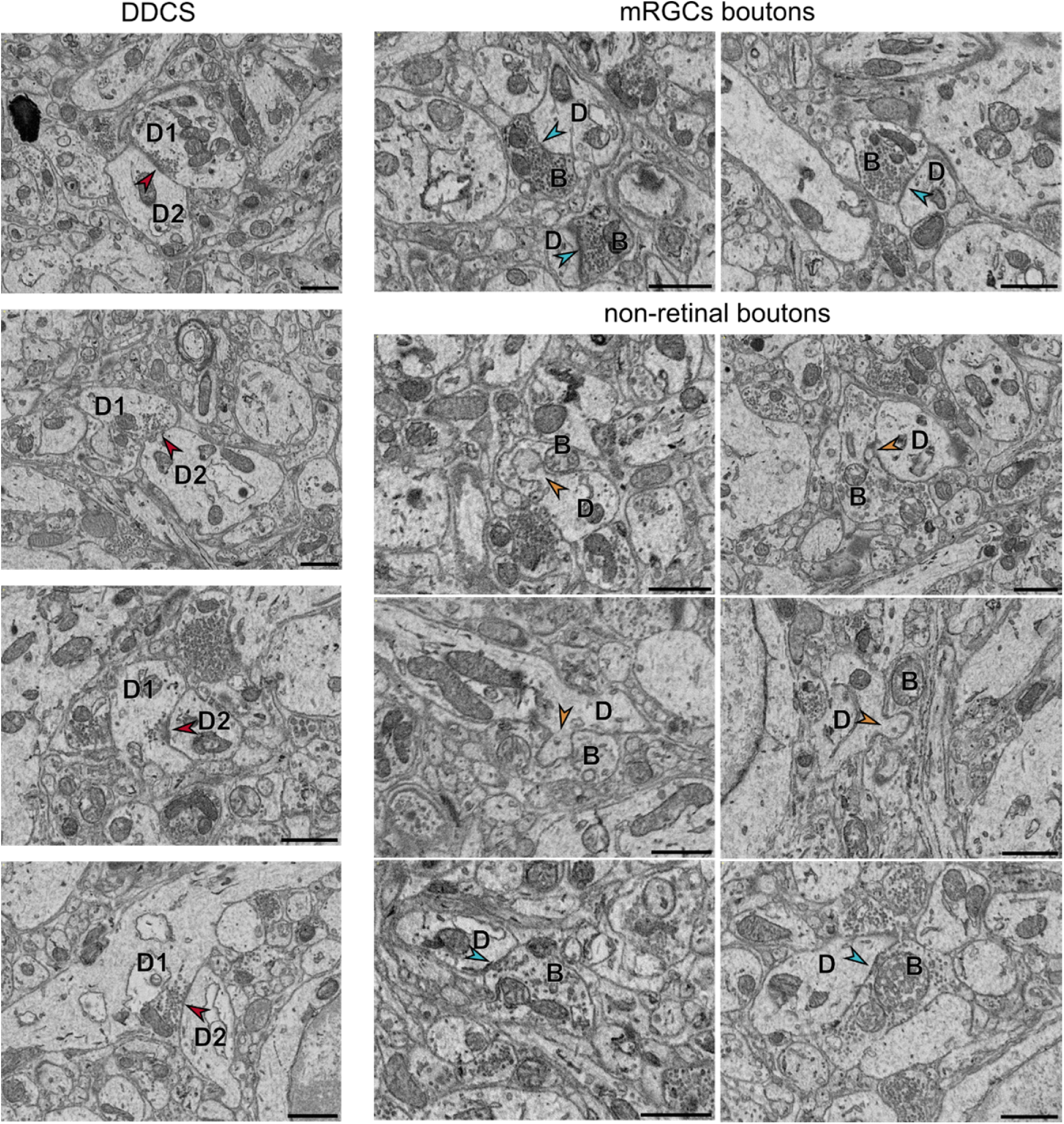
Examples of synapses in the SCN. Representative examples of EM images of DDCS (left) and boutons (right). Red arrowhead = DDCS; Blue arrowhead = ADCS; orange arrowhead = dendritic intrusion; D = dendrite; B = Bouton. Scalebar = 1µm

## Notes

### Competing Interest Statement

The authors have declared no competing interest.

